# *In vitro* activity and *in vivo* efficacy against non-tuberculous mycobacteria of SPR719, the active moiety of the novel oral benzimidazole prodrug SPR720

**DOI:** 10.1101/2022.12.08.519697

**Authors:** Nicole Cotroneo, Suzanne S. Stokes, Michael J. Pucci, Aileen Rubio, Kamal Hamed, Ian A. Critchley

## Abstract

Nontuberculous mycobacterial pulmonary disease (NTM-PD) is increasing globally, with *Mycobacterium avium* complex (MAC) accounting for 80% of cases and *Mycobacterium abscessus* among the next most prevalent pathogens. Current treatment necessitates long-term administration of poorly tolerated and modestly effective antibiotic combinations, highlighting the need for new oral agents. SPR719, the active moiety of the benzimidazole phosphate prodrug SPR720, inhibits the ATPase subunits of DNA gyrase B, a target with no human homolog and not exploited by current antibiotics. We demonstrated SPR719 activity against MAC (MIC_90_, 2 µg/mL) and *M. abscessus* clinical isolates (MIC_90_, 4 µg/mL), including those resistant to current standard-of-care (SOC) agents. *In vivo* efficacy of SPR720 was demonstrated against *M. avium* ATCC 700898 in a chronic C3HeB/FeJ murine model of pulmonary infection, both as a monotherapy and in combination with clarithromycin, ethambutol and rifabutin. SPR720 monotherapy exhibited a dose-dependent reduction in bacterial burden, with the largest reduction observed when combined with clarithromycin and ethambutol. Efficacy of SPR720 was also demonstrated against *M. abscessus* subspecies *abscessus* 1513, a virulent multidrug-resistant strain in a prolonged acute model of pulmonary infection in mice. SPR720 monotherapy exhibited a dose-dependent reduction in bacterial burden with further reductions when combined with SOC agents, clarithromycin and amikacin ± clofazimine. Taken together, the *in vitro* activity of SPR720 against common NTM pathogens in concert with the proof-of-concept efficacy in murine infection models warrants the continued clinical evaluation of SPR720 as a new oral option for the treatment of NTM-PD.

## INTRODUCTION

Nontuberculous mycobacterial pulmonary disease (NTM-PD) is a chronic progressive condition that is acquired via inhalation of nontuberculous mycobacteria (NTM) from environmental sources. The most common species causing pulmonary disease are members of the *Mycobacterium avium* complex (MAC) that have been implicated in approximately 80% of NTM-PD cases, including *M. avium, Mycobacterium intracellulare* and *Mycobacterium chimaera* (1-3). Most patients experience chronic cough, sputum production, shortness of breath, and, as the disease progresses, many report fatigue, night sweats, loss of appetite, and weight loss. NTM-PD due to MAC exhibits two major forms: a fibrocavitary form typically observed in middle-aged or older male smokers, which is a more severe disease that can rapidly damage lung tissue, and a nodular bronchiectatic form that occurs when infection develops in the small airways and air sacs resulting in chronic inflammation and bronchiectasis, usually observed in thin older females with no history of smoking (4, 5). In addition to MAC, rapidly growing species of the *Mycobacterium abscessus* complex are implicated in patients with pre-existing pulmonary disease resulting in progressive decline in pulmonary function and impaired quality of life (6).

Treatment of NTM-PD due to MAC typically involves a combination of different oral antibacterial agents taken over an extended period of 12 months after sputum conversion to culture negative (i.e., a total of approximately 15 to 18 months). First-line agents include macrolides (azithromycin or clarithromycin), ethambutol and rifamycins; these combinations exhibit variable efficacy, and many patients experience tolerability issues that often result in regimen modifications, thus highlighting the need for new classes of oral agents that may be included in new standard-of-care combination regimens (1). Treatment of pulmonary infections caused by *M. abscessus* complex with a combination of oral and parenteral antibiotics is complicated by multidrug resistance. Although clarithromycin, amikacin, and imipenem are current drugs of choice, new treatments are needed due to increasing intrinsic resistance to macrolides (7, 8).

SPR719, the active moiety of the phosphate prodrug SPR720 (Figure 1), belongs to a novel chemical class (aminobenzimidazoles) targeting the ATPase site located on the gyrase B subunit of the heterotetrameric bacterial gyrase protein, a mechanism that is distinct from that of fluoroquinolones. SPR719 has demonstrated antibacterial activity against NTM pathogens including MAC and *M. abscessus* clinical isolates (9, 10), has been shown to penetrate THP-1 monocytes which is critical for targeting these intracellular pathogens (11), and is currently being evaluated in a Phase 2 dose-ranging clinical study in patients with NTM-PD due to MAC (NCT05496374).

**FIG 1:**
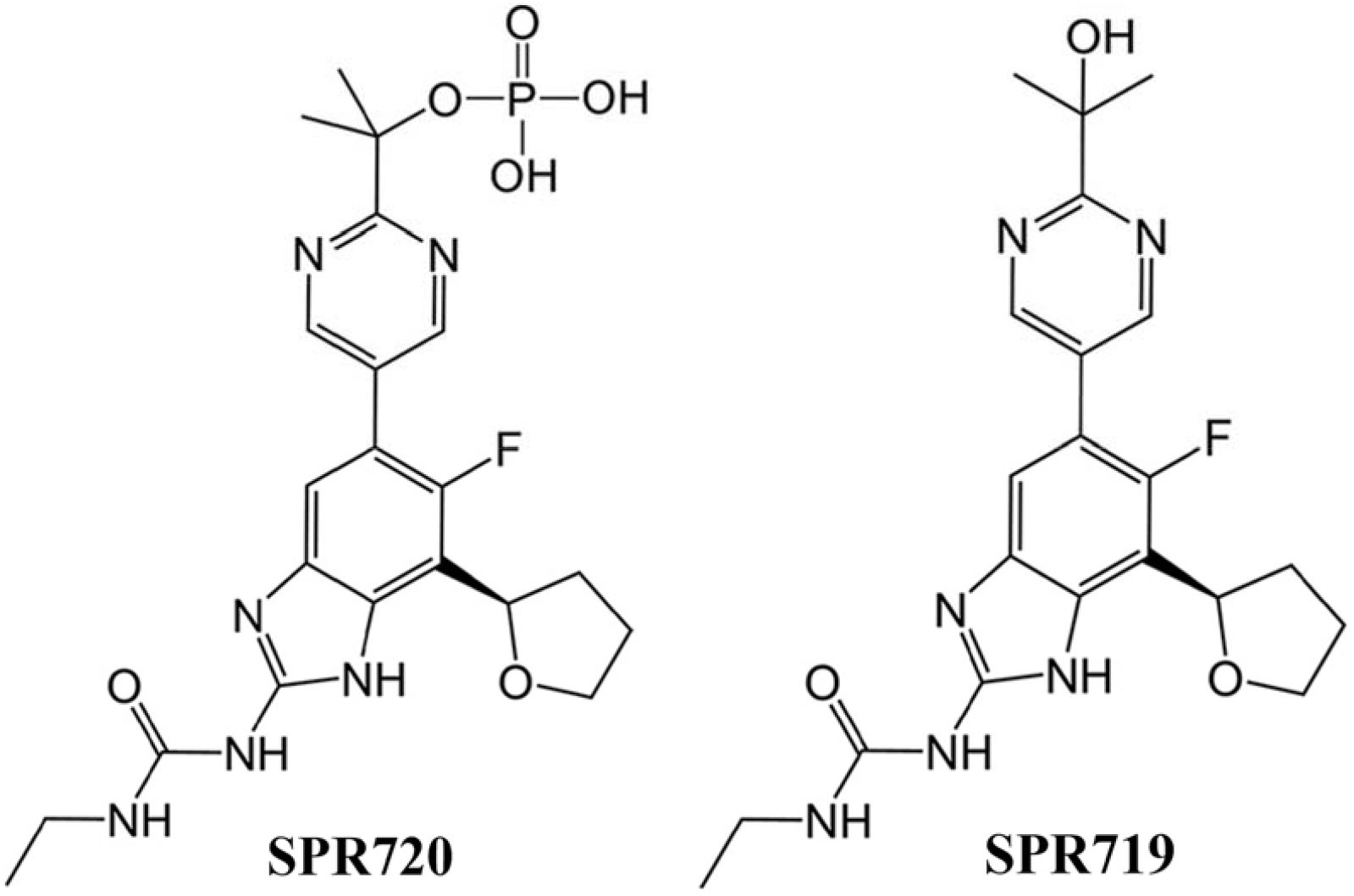
Chemical structure of SPR720 (oral phosphate prodrug) and SPR719 (active moiety)

The goals of the current study were to evaluate the *in vitro* antibacterial activity of SPR719 against clinical isolates of MAC and *M. abscessus* and to assess the *in vivo* efficacy alone and in combination with standard-of-care treatment regimens against *M. avium* and *M. abscessus* in murine infection models (12).

## RESULTS

*In vitro* studies were carried out using the active moiety, SPR719, following CLSI methodology with identification of resistant phenotypes as defined by CLSI criteria (13, 14) and *in vivo* studies employed the orally bioavailable phosphate prodrug, SPR720. Table 1 shows the susceptibility results for SPR719 and comparator agents against 105 clinical MAC isolates collected in the US and Japan from 2015 to 2018. All MAC isolates were inhibited by SPR719 at concentrations ≤4 µg/mL and the MIC_90_ value was 2 µg/mL. Among the clarithromycin-resistant isolates, the MICs for SPR719 ranged from 0.5 to 2 µg/mL and the MIC_90_ value was 2 µg/mL. The MICs for SPR719 against the 47 moxifloxacin-resistant MAC isolates ranged from 0.12 to 4 µg/mL and the MIC_90_ value was 2 µg/mL, confirming that resistance to fluoroquinolone agents does not impact susceptibility to SPR719, which was expected based on the different binding sites. There were 36 isolates that were resistant to ≥2 antibiotics with SPR719 MICs that ranged from 0.12 to 2 µg/mL.

**TABLE 1.** Anti-mycobacterial activity of SPR719 against clinical isolates of *M. avium* complex and *M. abscessus* collected from diverse specimen sources from patients in the US and Japan between 2015 and 2018

| Organism | Phenotype<sup>a</sup> | N | SPR719 MIC (µg/mL) |  |  |
| --- | --- | --- | --- | --- | --- |
|  |  |  | Range | MIC<sub>50</sub> | MIC<sub>90</sub> |
| *M. avium* complex | All | 105 | 0.002 – 4 | 1 | 2 |
|  | Amikacin-resistant<sup>b</sup> | 6 | 0.25 – 2 | NA | NA |
|  | Clarithromycin-resistant | 10 | 0.5 – 2 | 1 | 2 |
|  | Linezolid-resistant | 51 | 0.12 – 2 | 1 | 2 |
|  | Moxifloxacin-resistant | 47 | 0.12 – 4 | 1 | 2 |
|  | Resistant to ≥2 antibiotics | 36 | 0.12 – 2 | 1 | 2 |
| *M. abscessus* | All | 53 | 0.12 – 8 | 2 | 4 |
|  | Amikacin-resistant | 2 | 0.5 – 2 | NA | NA |
|  | Clarithromycin-resistant | 31 | 0.25 – 8 | 2 | 4 |
|  | Doxycycline-resistant | 41 | 0.12 – 8 | 2 | 4 |
|  | Cefoxitin-resistant | 2 | 2 – 4 | NA | NA |
|  | Imipenem-resistant | 9 | 0.5 – 2 | NA | NA |
|  | Linezolid-resistant | 6 | 1 – 4 | NA | NA |
|  | Moxifloxacin-resistant | 51 | 0.25 – 8 | 2 | 4 |
|  | Trimethoprim-sulfamethoxazole-resistant | 52 | 0.25 – 8 | 2 | 4 |

<sup>a</sup>Resistant phenotypes were defined according to CLSI interpretive criteria (CLSI M62 1<sup>st</sup> Edition)
<sup>b</sup>Amikacin resistance in MAC was determined using the breakpoints for the liposomal inhaled formulation

Susceptibility results for SPR719 against 53 clinical isolates of *M. abscessus* are also shown in Table 1 with all isolates being inhibited by SPR719 at concentrations ≤8 µg/mL with an MIC_90_ of 4 µg/mL, including isolates that were resistant to standard-of-care agents. There were 31 isolates that were resistant to clarithromycin with SPR719 MICs ranging from 0.25 to 8 µg/mL and 41 isolates that were doxycycline resistant with SPR719 MICs ranging from 0.12 to 8 µg/mL. As with MAC, resistance to moxifloxacin did not impact susceptibility to SPR719.

*M. avium* ATCC 700898, the organism evaluated for efficacy in the chronic C3HeB/FeJ murine infection model, was susceptible to SPR719 with an MIC of 2 µg/mL (Table 2). The MICs for clarithromycin, rifabutin and ethambutol were 1, 0.25 and 4 µg/mL, respectively. *M. abscessus* strain 1513 was the organism selected for evaluation of efficacy in the prolonged acute severe combined immune deficient (SCID) mouse model and was susceptible to SPR719 with an MIC of 2 µg/mL (Table 2). This organism was resistant to clarithromycin as well as other standard-of-care agents including ciprofloxacin, doxycycline, linezolid, sulfamethoxazole, and trimethoprim-sulfamethoxazole (Table 2).

**TABLE 2.** Activity of SPR719 and standard-of-care comparator agents against *M. avium* ATCC 700898 and *M. abscessus* 1513 strains evaluated in murine efficacy models

| *In vivo* test organism | Agent | MIC (µg/mL) | S/I/R interpretation<sup>a</sup> |
| --- | --- | --- | --- |
| *M. avium* ATCC 700898 | SPR719 | 2 | n/a |
|  | Clarithromycin | 1 | S |
|  | Rifabutin | 0.25 | n/a |
|  | Ethambutol | 4 | n/a |
| *M. abscessus* 1513 | SPR719 | 2 | n/a |
|  | Clarithromycin | >8 | R |
|  | Amikacin | 2 | S |
|  | Cefoxitin | 128 | R |
|  | Ciprofloxacin | >4 | R |
|  | Doxycycline | >16 | R |
|  | Linezolid | >16 | R |
|  | Sulfamethoxazole | >128 | n/a |
|  | Trimethoprim-sulfamethoxazole | >4/76 | R |

<sup>a</sup>Susceptibility was assessed according to CLSI interpretive criteria (CLSI M62 1<sup>st</sup> Edition)

Table 3 shows the changes in bacterial burden of *M. avium* ATCC 700898 in the lung, spleen, and liver of C3HeB/FeJ mice following treatment with different regimens of SPR720 alone and in combination with clarithromycin ± rifabutin, ethambutol or both. The bacterial burden in the day 61 untreated controls in the lung, spleen and liver were 6.30±.04, 4.20±0.04 and 5.10±0.08 log_10_ CFU/mL (log_10_ CFU/mL ± SEM), respectively. SPR720 alone showed dose-dependent reductions in bacterial burden in all tissues when compared with the day 61 untreated control. While SPR720 in combination with other standard-of-care agents showed varying degrees of reduction in bacterial burden, SPR720 in combination with clarithromycin and ethambutol showed the largest reduction in bacterial burden in the lung, spleen, and liver (Table 3). Figure 2 shows the changes in pulmonary bacterial burden (log_10_ CFU/lung) after treatment with SPR720 alone and in combination with other agents. SPR720 monotherapy was evaluated at 10, 30 and 100 mg/kg dosed once daily (QD) resulting in a dose-dependent reduction of *M. avium* ATCC 700898 at all doses tested when compared with the day 61 control group. Experimental groups of mice were also evaluated for lung histology and acid-fast bacilli on day 61. Mice receiving SPR720 at 30 mg/kg in combination with clarithromycin at 250 mg/kg and ethambutol at 100 mg/kg showed significantly reduced granuloma formation and prevalence of acid-fast bacilli as compared to untreated controls (Figure 3).

**TABLE 3.** *M. avium* ATCC 700898 bacterial burden (log_10_ CFU/organ) in the lung, spleen, and liver of C3HeBFeJ mice after treatment with oral SPR720 alone and in combination with clarithromycin ± rifabutin and/or ethambutol or both

| Treatment group regimens | No. of mice | Log<sub>10</sub> CFU ± SEM |  |  |
| --- | --- | --- | --- | --- |
|  |  | Lung | Spleen | Liver |
| Day 1 Pretreatment Control | 3 | 5.52±0.05 | 0±0 | 0±0 |
| Day 27 Pretreatment Control | 3 | 5.72±0.14 | 3.16±0.24 | 3.33±0.05 |
| Day 61 Control | 3 | 6.30±0.04 | 4.20±0.04 | 5.10±0.08 |
| SPR720 (10 mg/kg, QD) | 6 | 5.86±0.19 | 3.80±0.07 | 3.88±0.10 |
| SPR720 (30 mg/kg, QD) | 6 | 5.21±0.44 | 3.31±0.17 | 3.40±0.18 |
| SPR720 (100 mg/kg, QD) | 6 | 4.26±0.05 | 3.21±0.45 | 3.50±0.52 |
| CLR (250 mg/kg, QD) | 6 | 4.38±0.17 | 2.41±0.56 | 2.69±0.11 |
| CLR (250 mg/kg, QD) and RFB (100 mg/kg, QD) | 6 | 4.23±0.06 | 2.80±0.15 | 3.76±0.06 |
| CLR (250 mg/kg, QD) and EMB (100 mg/kg, QD) | 6 | 2.78±0.09 | 2.2±0.54 | 3.60±0.52 |
| CLR (250 mg/kg, QD), RFB (100 mg/kg, QD), and EMB (100 mg/kg, QD) and | 6 | 2.85±0.04 | 2.81±0.18 | 3.73±0.03 |
| CLR (250 mg/kg, QD) and SPR720 (30 mg/kg, QD) | 6 | 2.34±0.27 | 2.24±0.52 | 2.68±0.73 |
| CLR (250 mg/kg, QD), RFB (100 mg/kg, QD), and SPR720 (30 mg/kg, QD) | 6 | 2.50±0.27 | 2.40±0.64 | 2.73±0.57 |
| CLR (250 mg/kg, QD), EMB (100 mg/kg, QD) and SPR720 (30 mg/kg, QD) | 6 | 1.93±0.57 | 1.70±0.40 | 2.13±0.67 |
| CLR (250 mg/kg, QD), RFB (100 mg/kg, QD), EMB (100 mg/kg, QD) and SPR720 (30 mg/kg, QD) | 6 | 3.61±0.12 | 2.85±0.16 | 3.60±0.52 |

**FIG 2.**
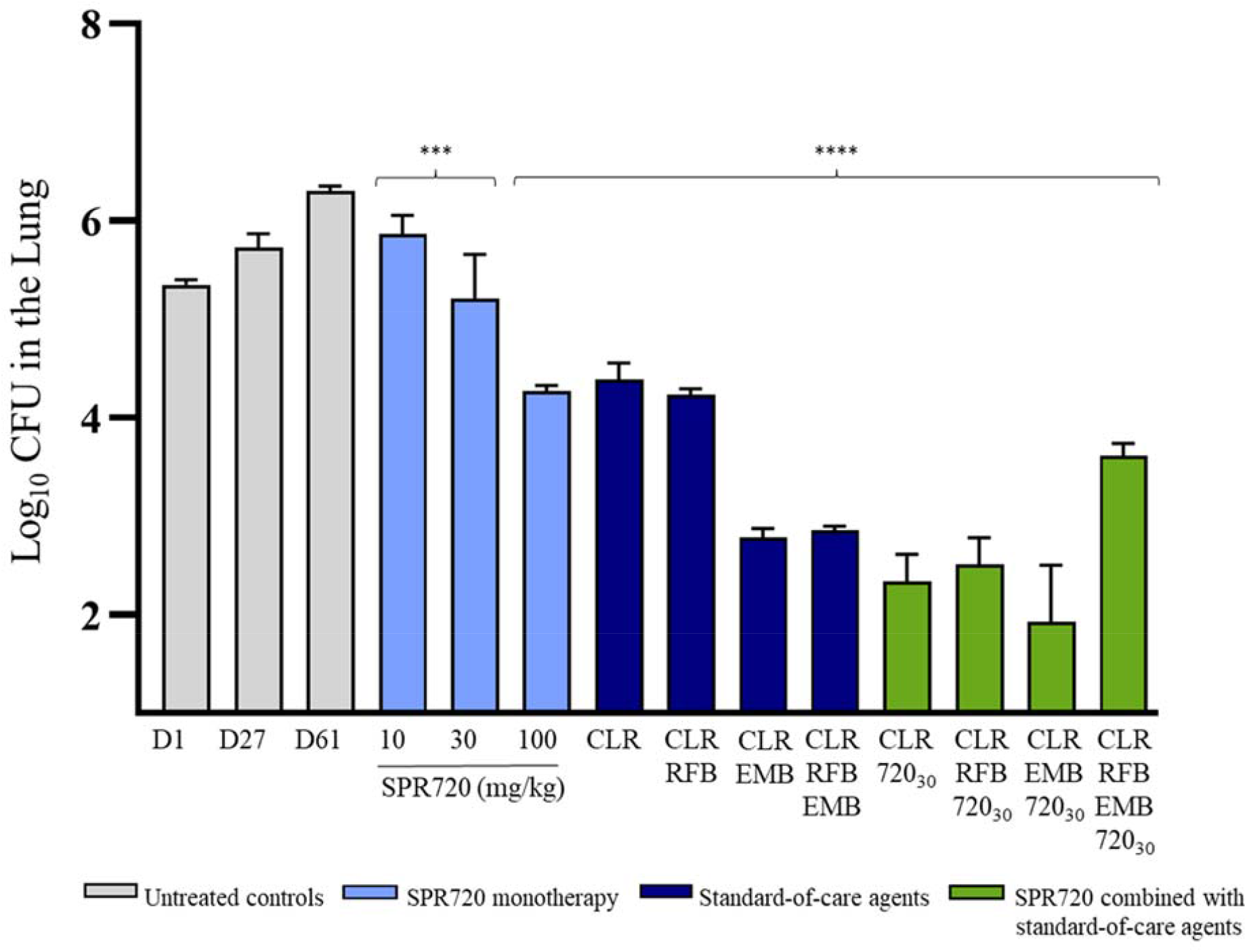
Pulmonary bacterial burden (log_10_ CFU/lung) of *M. avium* ATCC 700898 in C3HeBFeJ mice after treatment with oral SPR720 alone and in combination with clarithromycin ± rifabutin, ethambutol or both in a chronic murine infection model ***P value <0.001; ****P value <0.0001 relative to the day 61 untreated control. **Dosing regimens:** Clarithromycin (CLR), PO, QD at 250 mg/kg; Ethambutol (EMB), PO, QD at 100 mg/kg; Rifabutin (RFB), PO, QD at 100 mg/kg; SPR720, PO, QD at 10 mg/kg, 30 mg/kg and 100 mg/kg; SPR720_30_ indicates SPR720 was dosed at 30 mg/kg, PO, QD in combination with standard-of-care agents.

**FIG 3.**
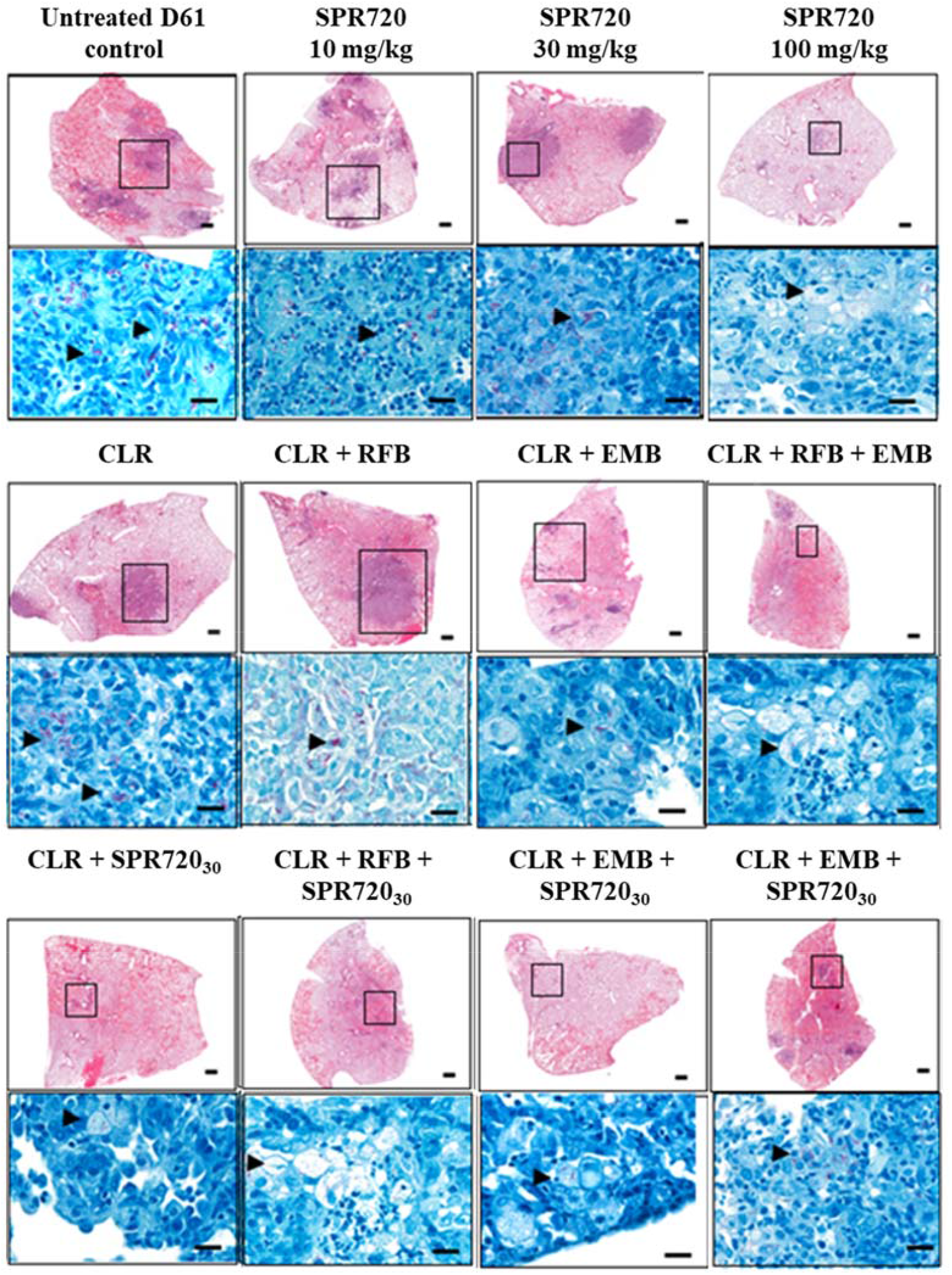
Pulmonary pathology in C3HeB/FeJ mice infected with *M. avium* ATCC 700898 after treatment with oral SPR720 alone and in combination with clarithromycin ± rifabutin, ethambutol or both. **Legend:** Clarithromycin (CLR), PO, QD at 250 mg/kg; Ethambutol (EMB), PO, QD at 100 mg/kg; Rifabutin (RFB), PO, QD at 100 mg/kg; SPR720, PO, QD at 10 mg/kg, 30 mg/kg and 100 mg/kg; SPR720_30_ indicates SPR720 was dosed at 30 mg/kg, PO, QD in combination with standard-of-care agents. Boxed areas in the top frames include the magnified area highlighted in the bottom frames; arrows denote the presence of acid-fast bacilli.

The reduction in bacterial burden achieved with all the combinations that included clarithromycin was statistically significant when compared with the day 61 control (p<0.0001). SPR720 at 30 mg/kg/day improved the efficacy of clarithromycin resulting in a 3.96 log_10_ CFU reduction in bacterial burden in the lung; the level of reduction of bacterial burden was similar to clarithromycin plus ethambutol, which resulted in a 3.52 log_10_ CFU reduction in bacterial burden in the lung when compared with the day 61 control (Table 3). A similar log reduction of 3.45 log_10_ CFU was observed with clarithromycin combined with ethambutol when combined with rifabutin (100 mg/kg). The greatest reduction in bacterial burden was observed with the combination of clarithromycin (250 mg/kg), ethambutol (100 mg/kg) and SPR720 (30 mg/kg) with an average 4.37 log_10_ CFU reduction bacterial burden in the lung, while bacterial burden in the lung was reduced by only 2.70 log_10_ CFU relative to the day 61 untreated control when rifabutin (100 mg/kg) was added to this regimen.

Table 4 shows the changes in bacterial burden of *M. abscessus* 1513 in the lung, spleen and liver of mice following different treatment regimens of SPR720 alone and in combination with standard-of-care agents. Untreated mice were evaluated on days 1, 27 and 61 post-infection to assess bacterial growth in the lung liver and spleen. The bacterial burden in the day 61 untreated controls in the lung, spleen and liver was 6.55±0.08, 6.74±0.05 and 7.22±0.02 log_10_ CFU, respectively. SPR720 monotherapy showed a dose-dependent reduction in bacterial burden after 28 days of treatment in all tissues as compared to the day 61 untreated control. SPR720 in combination with other standard-of-care regimens also showed varying degrees of reduction. The greatest reduction in bacterial burden in all tissues occurred in mice treated with SPR720 (50 mg/kg) in combination with clarithromycin (250 mg/kg), amikacin (150 mg/kg) and clofazimine (50 mg/kg) QD. The changes in bacterial burden in the lung for SPR720 alone and in combination are shown as a histogram in Figure 4 that highlights the dose-dependent reduction in log_10_ CFU in the lung with SPR720 monotherapy at 25, 50 and 100 mg/kg QD.

**TABLE 4.** *M. abscessus* subspecies *abscessus* 1513 bacterial burden (log_10_ CFU/organ) in the lung, spleen and liver of SCID mice after treatment with oral SPR720 alone and in combination with clarithromycin and amikacin ± clofazimine

| Treatment group regimens | No. of mice | Log<sub>10</sub>CFU ± SEM |  |  |
| --- | --- | --- | --- | --- |
|  |  | Lung | Spleen | Liver |
| Day 1 Pretreatment Control | 3 | 5.83±0.04 | 5.58±0.04 | 6.12±0.04 |
| Day 27 Pretreatment Control | 6 | 6.39±0.03 | 5.77±0.11 | 7.03±0.11 |
| Day 61 Untreated Control | 6 | 6.55±0.08 | 6.74±0.04 | 7.22±0.02 |
| SPR720 (25 mg/kg QD) | 6 | 4.45±0.07 | 4.25±0.20 | 5.07±0.16 |
| SPR720 (50 mg/kg QD) | 6 | 3.50±0.22 | 3.89±0.18 | 4.91±0.29 |
| SPR720 (100 mg/kg QD) | 6 | 2.71±0.40 | 3.30±0.16 | 4.73±0.41 |
| CFZ (20 mg/kg QD) | 6 | 5.08±0.05 | 4.63±0.13 | 5.83±0.04 |
| AMK (150 mg/kg QD) | 6 | 4.53±0.08 | 4.09±0.04 | 5.23±0.07 |
| CLR (250 mg/kg QD) | 6 | 4.46±0.17 | 5.06±0.05 | 5.45±0.05 |
| CLR (250 mg/kg QD) and AMK (150 mg/kg QD) | 6 | 3.99±0.14 | 5.07±0.05 | 4.86±0.12 |
| CLR (250 mg/kg), AMK (150 mg/kg) and CFZ (20 mg/kg QD) | 6 | 3.70±0.25 | 4.83±0.11 | 4.33±0.07 |
| CLR (250 mg/kg QD) AMK (150 mg/kg QD) and SPR720 (50 mg/kg QD) | 6 | 2.20±0.22 | 3.93±0.11 | 4.92±0.19 |
| CLR (250 mg/kg QD), AMK (150 mg/kg QD), CFZ (20 mg/kg QD) and SPR720 (50 mg/kg QD) | 6 | 1.86±0.16 | 3.36±0.21 | 3.96±0.13 |

**FIG 4:**
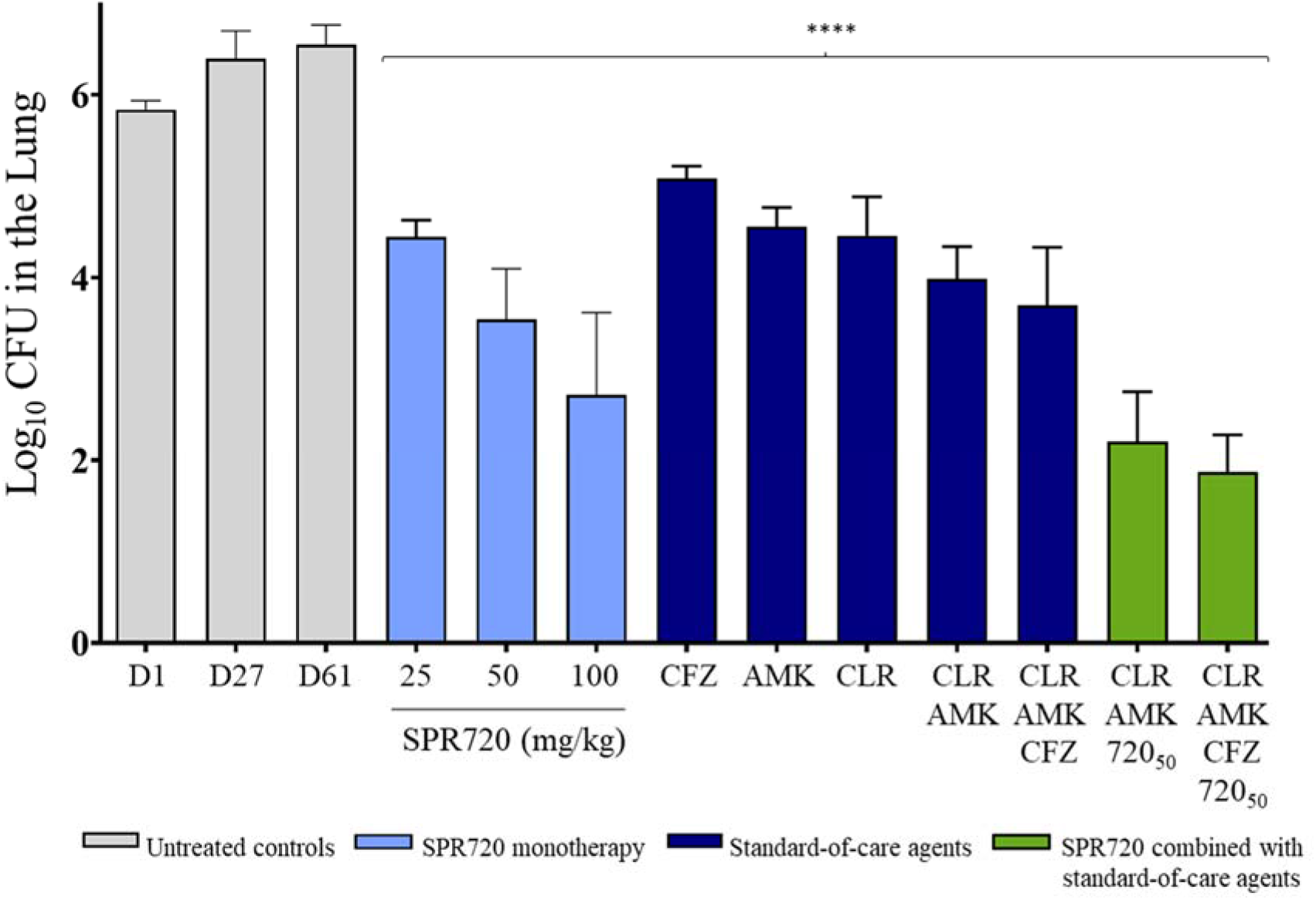
Pulmonary bacterial burden (log_10_ CFU/lung) of MDR *M. abscessus* subspecies *abscessus* in SCID mice after treatment with oral SPR720 alone and in combination with clarithromycin, amikacin and clofazimine in a proloned acute murine infection model ****P value <0.0001 relative to the day 61 untreated control group. **Dosing regimens:** Amikacin (AMK), subcutaneously (SC), QD at 150 mg/kg; Clarithromycin (CLR), PO, QD at 250 mg/kg; Clofazimine (CFZ), PO, QD at 20 mg/kg; SPR720, PO, QD at 25 mg/kg, 50 mg/kg and 100 mg/kg; SPR720_50_ indicates SPR720 was dosed at 50 mg/kg, PO, QD in combination with standard-of-care agents.

## DISCUSSION

The burden of NTM-PD continues to increase in the United States and globally (15, 16) and is now widely recognized as an important cause of morbidity and mortality. Combination therapy has become the mainstay with a macrolide in combination with ethambutol and a rifamycin being considered the standard-of-care regimen for NTM-PD due to MAC and a macrolide with one or more parenteral agents (amikacin, cefoxitin or imipenem) for NTM-PD due to *M. abscessus* (4). Current therapeutics remain challenging because of the long treatment duration and poor tolerability, with many patients left with residual lung dysfunction (15, 17) or failing to respond to treatment, relapsing, or developing macrolide resistance (18).

SPR720 is a novel aminobenzimidazole antimicrobial phosphate prodrug that prevents bacterial growth by inhibiting bacterial DNA gyrase (19, 20) and is orally bioavailable in humans (21). Specifically, SPR719 targets ATPase located on the bacterial gyrase B subunit of the heterotetrameric gyrase protein, a mechanism that is distinct from fluoroquinolones and other currently marketed antibiotics, so it is unlikely for SPR719 to show cross-resistance with any standard-of-care agents due to alteration of the target enzyme. Furthermore, SPR719 exhibits antibacterial activity against many commonly encountered NTM pathogens including *M. avium, M. intracellulare, M. abscessus*, and other important species (9); the results of the susceptibility testing in this study confirmed that SPR719 exhibited consistent activity against both MAC and *M. abscessus* isolates that were resistant to standard-of-care agents, including those which are moxifloxacin resistant. Laboratory studies have identified spontaneous mutants of *M. avium* resistant to SPR719 at a frequency of 10^−8^ CFU at 2x MIC with a Ile173Thr mutation in the ATPase domain of gyrase B; resistant mutants did not exhibit cross-resistance to clarithromycin or moxifloxacin (22). SPR720 is anticipated to be used clinically in combination therapy to mitigate such resistance development that would be unavoidable with any monotherapy. In time-kill assays, SPR719, in combination with ethambutol, increased bacterial killing and suppressed outgrowth of *M. avium* (10). Similarly, in this study, SPR720 in combination with clarithromycin and ethambutol showed the greatest reduction in bacterial burden of *M. avium* in the chronic murine lung model while the inclusion of rifabutin in this regimen reduced overall efficacy. Although the precise mode of action of ethambutol has not been established, it is thought to inhibit cell wall biosynthesis (arabinogalactan) in mycobacteria (23) and may act synergistically to enhance access of SPR719 to intracellular targets in MAC. These results suggest that SPR720 could be a potential substitute for the rifamycins in the current standard-of-care regimens, which would be an advantage since rifamycins are potent inducers of cytochrome P450 enzymes and transporters, and drug-drug interactions have been reported during tuberculosis treatments (24). Patients treated with rifabutin commonly experience serious side effects including, but not limited to, arthralgias, uveitis, leukopenia, thrombocytopenia, “flu-like syndrome,” and elevated hepatic transaminases and should be monitored closely for rifabutin-related toxicity (25, 26). The replacement of rifabutin as a standard-of-care therapeutic with a more well-tolerated antibiotic may significantly reduce morbidity among patients who experience these side effects. Potential limitations of this *M. avium* murine efficacy study include that dosing regimens of SPR720 doses >100 mg/kg were not explored to identify the maximal effect and the *M. avium* strain ATCC 700898 evaluated was susceptible to clarithromycin.

The *M. abscessus* strain 1513 that was evaluated in the prolonged acute SCID treatment mouse model was highly virulent and resistant to clarithromycin and other standard-of-care agents. *M. abscessus* pathogens are formidable and difficult to treat, most notably due to the presence of an inducible erythromycin methylase (*erm*) gene that confers resistance to macrolides (7). The dose-dependent reduction in pulmonary bacterial burden observed with SPR720 monotherapy in this murine model demonstrates the efficacy against an organism that is resistant to clarithromycin and many other standard-of-care agents. Furthermore, the addition of SPR720 to the standard-of-care agents such as clarithromycin, amikacin and clofazimine resulted in a greater reduction in bacterial burden in the lung when compared with the standard-of-care agents.

Collectively, the results of this study highlight the antibacterial activity of SPR719 against commonly encountered NTM organisms, including those resistant to current antibiotics. SPR720 has demonstrated proof-of-concept efficacy by exhibiting dose response as a stand-alone agent and even greater reductions in bacterial burden when evaluated in combination with other standard-of-care agents. These results support the ongoing clinical development of SPR720 as a new agent for the treatment of NTM-PD.

## MATERIALS AND METHODS

### *In Vitro* Susceptibility testing

Clinical isolates of MAC and *M. abscessus* were tested for susceptibility to SPR719 in accordance with Clinical and Laboratory Standard Institute (CLSI) guidelines (23)(13). *M. abscessus* was tested in cation-adjusted Mueller-Hinton broth (CAMHB) while *M. avium* was tested in CAMHB supplemented with 5% oleic acid albumin dextrose catalase complex (OADC). In the susceptibility profiling study, 105 clinical MAC isolates were collected from diverse specimen sources from patients in different geographic in the United States and Japan between 2015 and 2018 were tested for susceptibility to SPR719 and included isolates that were susceptible to standard-of-care agents and isolates with resistant phenotypes as defined by CLSI criteria (14). A total of 53 clinical isolates of *M. abscessus* were also tested for susceptibility to SPR719 including isolates that were susceptible and resistant to standard-of-care agents.

### Organisms evaluated in murine efficacy models

*M. avium* (Chester) ATCC 700898 is a standard strain widely used for MAC susceptibility testing and selected for evaluation in the chronic C3HeBFeJ mouse infection model. The isolate was tested for susceptibility to the active moiety SPR719 and standard-of-care comparator agents including clarithromycin, ethambutol and rifabutin that were also evaluated for efficacy in the mouse infection model. *M. abscessus* subspecies *abscessus* strain 1513 was evaluated in the prolonged acute SCID murine model of pulmonary and systemic infection and tested for susceptibility to SPR719 and standard-of-care comparator agents. All procedures involving animals were approved by the Colorado State University Animal Care and Use Committee.

### Chronic C3HeBFeJ mouse infection model

A 60-day chronic C3HeB/FeJ mouse infection model was developed to be representative of NTM-PD since these mice form foci of necrosis in granulomas, as previously described (12). Mice were infected by aerosol delivery of 1 × 10^8.5^ CFU/mL of *M. avium* ATCC 700898 that was susceptible to SPR719 with an MIC of 2 µg/mL. Untreated groups of mice were evaluated on days 2, 27 and 61 post-infection to assess bacterial growth in the lung, spleen and liver which were extracted, homogenized in 4.5 mL of phosphate buffered saline (PBS), plated on 7H11 agar plates, and incubated for ∼30 days to determine growth. On day 28 post-infection, the oral prodrug SPR720 was administered by oral gavage at 10 mg/kg/day, 30 mg/kg/day, and 100 mg/kg/day QD. Positive control clarithromycin was administered by oral gavage at 250 mg/kg/day. Ethambutol and rifabutin were also delivered by oral gavage at 100 mg/kg/day in combination treatments. The animals were dosed consecutively from day 28 to 60. At the end of the dosing phase (day 61), lung, spleen and liver were removed, processed, and plated on Middlebrook 7H11 media for ∼30 days to enumerate CFUs. Histological analysis was performed by removing one whole lung lobe from each mouse which was fixed with 10% formalin in PBS. Sections from these tissues were stained with haematoxylin-eosin and with Ziehl-Neelsen acid-fast stains (27, 28).

### Prolonged Acute SCID Treatment Mouse Model

An acute SCID model previously described by Obregon-Henao *et al*., (27) was used to evaluate the efficacy of SPR720 against *M. abscessus*. SCID Mice were infected with 1 × 10^6^ CFU/mouse via the tail vein with *M. abscessus* subspecies *abscessus* strain 1513 that was susceptible to SPR719 with an MIC of 2 µg/mL. Untreated groups of mice were evaluated on days 1, 27 and 61 post-infection to assess bacterial growth in the lung, spleen and liver. On day 28 post infection, SPR720 was administered as monotherapy by oral gavage at 25 mg/kg, 50 mg/kg and 100 mg/kg QD. Standard-of-care agents evaluated as monotherapy by oral gavage included clarithromycin (250 mg/kg, QD), amikacin (150 mg/g, QD) and clofazimine (20 mg/kg, QD). Treatment initiated 28 days post infection and continued until day 60. Bacterial burden in the lung, spleen and livers of infected mice were assessed on days 2, 21 and 61 after infection by plating onto Middlebrook 7H11 agar and enumerating CFUs after 7 days incubation at 37°C.

### Statistical analysis

Data are presented using the mean values from 3 to 6 mice per group performed in a single experiment. Statistical analysis was evaluated by a one-way ANOVA followed by a multiple comparison analysis of variance by a one-way Tukey test (SigmaStat software program). Data are presented using the mean values plus or minus the standard error of the mean (SEM). Significance was considered with a *P*-value of <0.05 (26).

## ACKNOWLEDGEMENTS

We thank Diane J. Ordway and Deepshikha Verma of the Colorado State University, Mycobacterial Research Laboratory, Department of Microbiology, Immunology, and Pathology (Fort Collins, CO, USA) for their assistance in the conduct of murine efficacy models described in the manuscript.

